# Neural representation of goal direction in the monarch butterfly brain

**DOI:** 10.1101/2022.10.15.512348

**Authors:** M. Jerome Beetz, Christian Kraus, Basil el Jundi

## Abstract

Neural processing of a navigational goal requires the continuous comparison between the current heading and the intended goal direction. While the neural basis underlying the current heading is well-studied in insects, the coding of the goal direction is completely unexplored. Here, we identify for the first time neurons that encode goal direction in the brain of a navigating insect, the monarch butterfly. The spatial tuning of these neurons accurately correlates with the animal’s goal direction while being unaffected by compass perturbations. Thus, they specifically encode the goal direction similar to goal neurons described in the mammalian brain. Taken together, a navigation network based on goal-direction and heading-direction neurons generates steering commands that efficiently guides the monarch butterflies to their migratory goal.

## INTRODUCTION

For goal-directed navigation, animals need to register their current orientation in space, as well as the direction of their goal. Consequently, their brain constantly compares the current heading direction with the goal direction (Dacke and el Jundi, 2018; Honkanen et al., 2019). While the former is encoded by evolutionarily conserved head-direction (HD) neurons found in different species (Beetz et al., 2022; Ben-Yishay et al., 2021; Geva-Sagiv et al., 2015; Hulse and Jayaraman, 2020; Petrucco et al., 2022; Seelig and Jayaraman, 2015; Takahashi et al., 2022; Taube et al., 1990; Varga and Ritzmann, 2016; Vinepinsky et al., 2020), goal-direction (GD) neurons whose action potential rate correlates with the animal’s goal direction have only been reported in the mammalian brain (Sarel et al., 2017). However, even in the tiny brain of an insect, a robust representation of the goal direction is of the highest ecological importance. For instance, monarch butterflies are well known for their spectacular southward migration over ~5,000 km from the Northern US and Canada to their overwintering site in Central Mexico. To maintain a goal direction during migration, the butterflies use a sun compass for orientation (Mouritsen and Frost, 2002), which is processed in a brain region termed the central complex (Heinze et al., 2013; Heinze and Reppert, 2011). Previous studies in a variety of insects have shown that the central complex houses HD neurons and steering neurons responsible for the animal’s steering behavior (Beetz *et al*., 2022; Martin et al., 2015; Seelig and Jayaraman, 2015; Varga and Ritzmann, 2016). Although a number of theoretical models predict that the central complex also houses GD neurons (Honkanen *et al*., 2019; Matheson et al., 2022; Stone et al., 2017), similar to the ones described in the bat hippocampus (Sarel *et al*., 2017), their existence to date has been completely speculative.

## RESULTS

We tethered monarch butterflies at the center of a flight simulator, in which they could freely steer in any goal direction with respect to a virtual sun (Fig. 1A). Although the tested butterflies were not in their migratory phase, they reliably maintained consistent goal directions (Fig. 1B; fig. S1). This goal-directed behavior likely emerges by matching the current heading - encoded by the butterflies’ compass - with an internal goal representation. To dissociate between these directional representations, we perturbed the butterflies’ compass without affecting their goal representation (Fig. 1C). This was achieved by displacing the sun along the azimuth every 90 s (fig. S2). To maintain the initial goal direction relative to the sun, the butterflies adjusted their heading direction in accordance with the new sun position (Fig. 1D). The behavioral response was independent of the size of sun displacement (fig. S3) and could be reliably evoked in all tested butterflies (Fig. 1E). Taken together we successfully shifted the polarity of the butterflies’ compass, while the animal’s goal direction remained unaffected. Thus, HD neurons in the butterfly central complex should change their spatial tuning, following compass perturbations, while the spatial tuning of GD neurons should remain invariant.

**Fig. 1.**
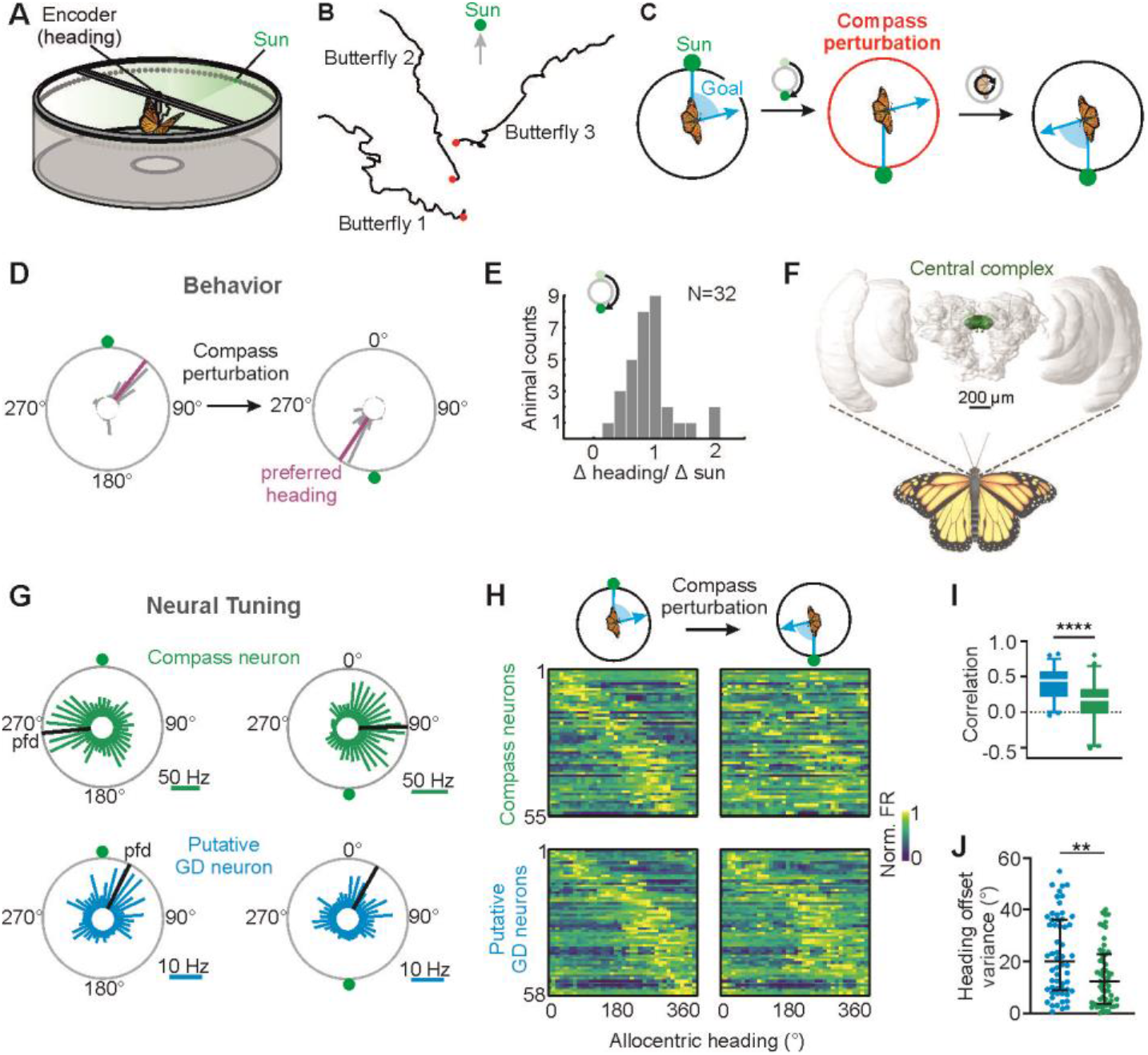
Monarch butterflies maintain goal directions relative to a virtual sun. (**A)** The arena’s inner circumference was equipped with green lights, allowing us to present a virtual sun and change its position. **(B)**Virtual flight trajectories (8-minute flights) of three tested butterflies. (**C**) Schematic drawing of the compass perturbation experiment through sun displacements. The compass polarity was manipulated while the butterfly’s goal direction remained unaffected (**D**) Change in heading direction of a butterfly after compass perturbation (180° sun displacement). (**E**) Butterflies changed their heading in accordance with the change in the position of the virtual sun. (**F**) Frontal view of the monarch butterfly brain with the central complex highlighted. (**G**) Tuning of two neurons prior to (*left*) and after (*right*) a 180° sun displacement. Black bars indicate the preferred firing directions (*pfds*). (**H**) Angular tuning of compass and putative goal direction (GD) neurons prior to (*left*) and after compass perturbation (*right*). Neurons are ordered according to their pfds before compass perturbation. (**I**) Correlation of the angular tuning prior to and after compass perturbation and (**J**) heading offsets variances in response to compass perturbations for putative GD (*blue*, n = 58) and compass (*green*, n = 55) neurons. Low heading offset variances indicate that the pfds were yoked to the butterfly’s heading.

While perturbing the butterflies’ compass system, we simultaneously monitored the neural activity of spatially tuned central-complex neurons (Fig. 1F, fig. S4). With tetrodes implanted in the central complex, we recorded from 113 neurons (~ 4.6 ± 2.2 neurons/animal) that showed a spatial tuning when the butterflies oriented in darkness (fig. S5), an important requirement for an internal representation of heading and goal directions (Beetz *et al*., 2022; Nyberg et al., 2022; Seelig and Jayaraman, 2015). As expected for compass neurons, i.e., HD neurons, we found neurons that substantially changed their angular tuning following compass perturbations (Fig. 1G, fig S6A). In total, 55 of 113 neurons (48.7%) modified the direction of their angular tuning, reflected by the preferred firing direction (pfd), according to the change in the animals’ heading (Fig. 1H). Variations in the action potential rate during flight could not explain these tuning shifts (p = 0.75, U = 1540; MWU, fig. S7). Thus, the tuning of these neurons reflects the butterfly compass system, similar to the *Drosophila* HD neurons (Green et al., 2019). Importantly, the angular tuning of another 58 neurons (51.3%) was unaffected by compass perturbations (Fig. 1G and 1H, fig S6A-S6C). The correlation between their angular tuning measured before and after compass perturbations was much higher than in compass neurons (p < 0.001, R^2^ = 0.18, unpaired t-test, Fig. 1I). Moreover, their tuning showed a higher variance of heading offsets (p = 0.005, U = 1113, MWU, Fig. 1J) indicating that they were *not* linked to the coding of the butterflies’ compass. Given that the animals’ goal direction remained consistent throughout compass perturbations (Fig. 1C), we hypothesized that these neurons encode the butterflies’ internal goal direction.

Conversely, the putative GD neurons could encode any stable cue in the environment, e.g., magnetic information (Wan et al., 2021). To exclude this possibility and ultimately test for goal coding, we next reset the butterflies’ goal direction - following compass perturbations - by applying small electric shocks to their necks whenever they headed towards their initial goal direction (Fig. 2A). This aversive conditioning did indeed reliably change the butterflies’ goal direction (129.7°± 39.9°; Fig. 2B and 2C; fig. S8A and S8B). Electric stimulation per se did not affect the orientation performance indicated by similarly high flight precision prior to and after conditioning (p = 0.63, R^2^ = 0.015, N = 17, paired t-test, fig. S9). Potential effects of the electric stimulations on neural tuning were excluded through control experiments (p = 0.63, W = 1136, n = 256, WSRT, fig. S8C). To ensure that we recorded from the same neurons throughout conditioning, we correlated the spike shapes within the neurons and across different periods and compared them with spike shapes across different neurons (fig. S10).

**Fig. 2.**
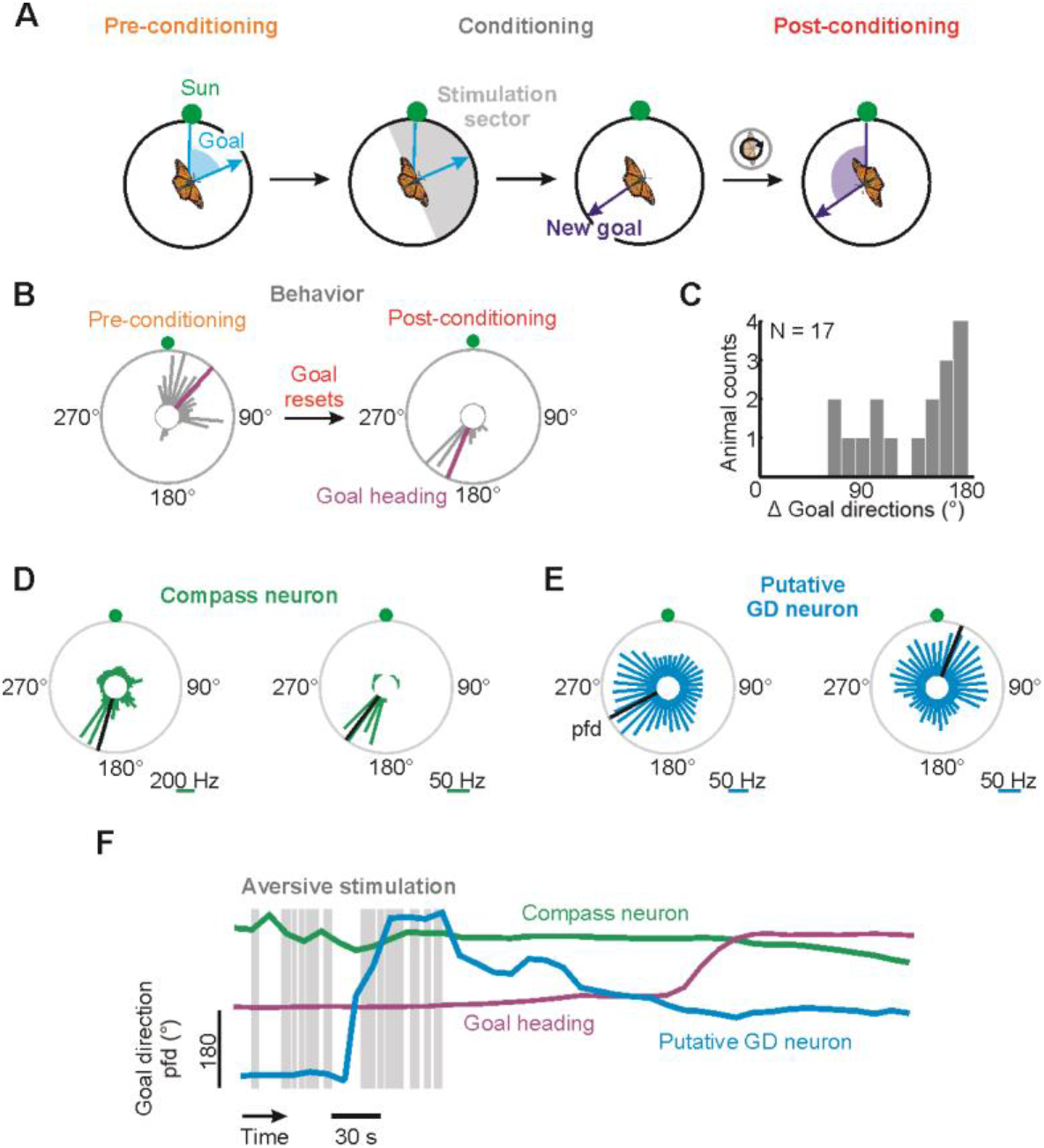
Resetting the goal direction. (**A**) We reset the goal direction by applying electric shocks to the butterflies’ neck whenever they set their initial goal direction (± 90°; stimulation sector). (**B**) Circular plots showing the heading before and after conditioning. Magenta lines indicate the Goal heading. (**C**) Changes in goal directions in 17 animals induced by aversive conditioning. (**D, E**) Tuning of a compass (D) and a putative goal-direction (E) neuron prior to (*left*) and after (*right*) resetting the goal direction. Black lines indicate the preferred firing directions (*pfds*). (**F**) Goal heading (magenta line) pfds of a compass (*green*) and a goal-direction (*blue*) neuron plotted as a function of time. Gray boxes highlight periods of electric stimulation.

If GD neurons exist in the insect central complex, we expected that their pfds should be tightly linked to butterflies’ new goal direction. Remarkably, in addition to compass neurons that did not change their angular tuning (Fig. 2D), we found neurons whose angular tuning changed in association with the butterflies’ goal directions (Fig. 2E and 2F). The neurons with pfds yoked to butterfly’s flight behavior observed during aversive conditioning are likely the GD neurons (fig. S11). Similar to GD neurons in mammals, the neural activity of GD neurons in butterflies should not represent the animals’ compass directions (Sarel *et al*., 2017). Therefore, we expected that the angular tuning of GD neurons should only change during aversive conditioning but *not* after compass perturbations (Fig. 3A). Interestingly, 20 neurons (31 %) exclusively shifted their pfds during aversive conditioning but showed invariant pfds during compass perturbations (Fig. 3B, *upper heatmaps*, p = 0.012, W = 132, n = 20, WSRT, Fig. 3C). In contrast, the angular tuning of 13 neurons (20 %) changed only when we perturbed the compass (Fig. 3B, *lower heatmaps*), clearly showing that these are HD neurons. Thus, while the angular tuning of HD neurons was specifically modulated during compass perturbations (p = 0.01, t = 2.72, unpaired t-test), the pfds of the GD neurons were only affected when the butterflies set a new goal direction (Fig. 3D, p < 10^-5^, t = 5.89, unpaired t-test). In addition, the pfds of the GD neurons were tightly linked to the goal direction, represented by relatively constant goal offsets (p < 10^-5^, U = 60, n = 39 GD & 13 HD neurons, MWT; Fig. 3E).

**Fig. 3.**
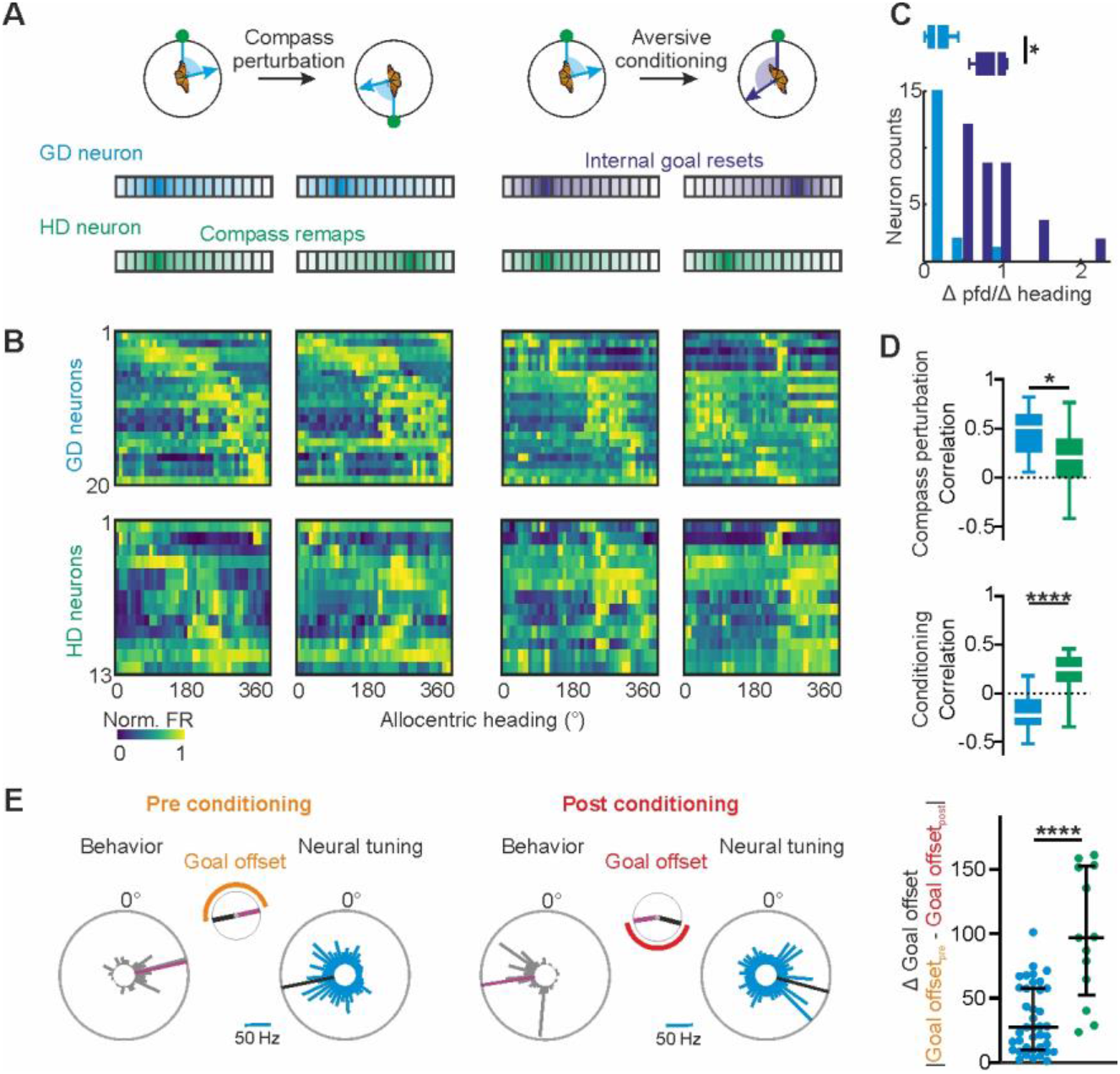
Goal coding in monarch butterflies. (**A**) Hypothesized neural tuning in response to compass perturbations and aversive conditioning for GD (*blue*) and HD (*green*) neurons. (**B**) Change in angular tuning of GD (upper row) and HD (lower row) neurons in responses to compass perturbations (two left columns) and aversive conditioning (two right columns). Neurons are ordered according to their pfds before compass perturbation and conditioning. (**C**) Ratio of changes in preferred firing directions (pfds) and heading changes for GD neurons during compass perturbations (bright blue data) and during conditioning (dark blue data). X-values close to 0 indicate no correlation between angular tuning and heading. (**D**) Correlation of angular tuning before and after compass perturbations (*top*) or conditioning (*bottom*). (**E**) Differences in goal offsets prior to (*left*) and after (right) conditioning in GD (*blue*) and HD (*green*) neurons.

Central-complex models predict that GD neurons are presynaptic to steering neurons that generate pre-motor steering commands. Hence, the tuning of GD and steering neurons should be closely associated (Matheson *et al*., 2022; Wystrach et al., 2020). During our experiments, we recorded from 19 neurons that showed tuning characteristics expected from steering cells. Their angular tuning was tightly linked to the butterflies’ change in flight direction during compass perturbation *and* aversive conditioning (Fig. 4A and 4B, fig S12). As typical for steering cells (Martin *et al*., 2015), the neurons modulated their firing rate prior to each turn of the animal (Fig. 4C). Interestingly, the GD neurons also increased their firing rates prior to flight turns (Fig. 4D). While GD neurons encoded equally strong left and right turns, steering neurons typically exhibited a directional selectivity to one rotation direction (Fig. 4E). Interestingly, GD neurons monitored simultaneously with steering neurons encoded turns even prior to the steering neurons (p = 0.005, W = −85, WSRT, Fig. 4F). This observation fits well with

**Fig. 4.**
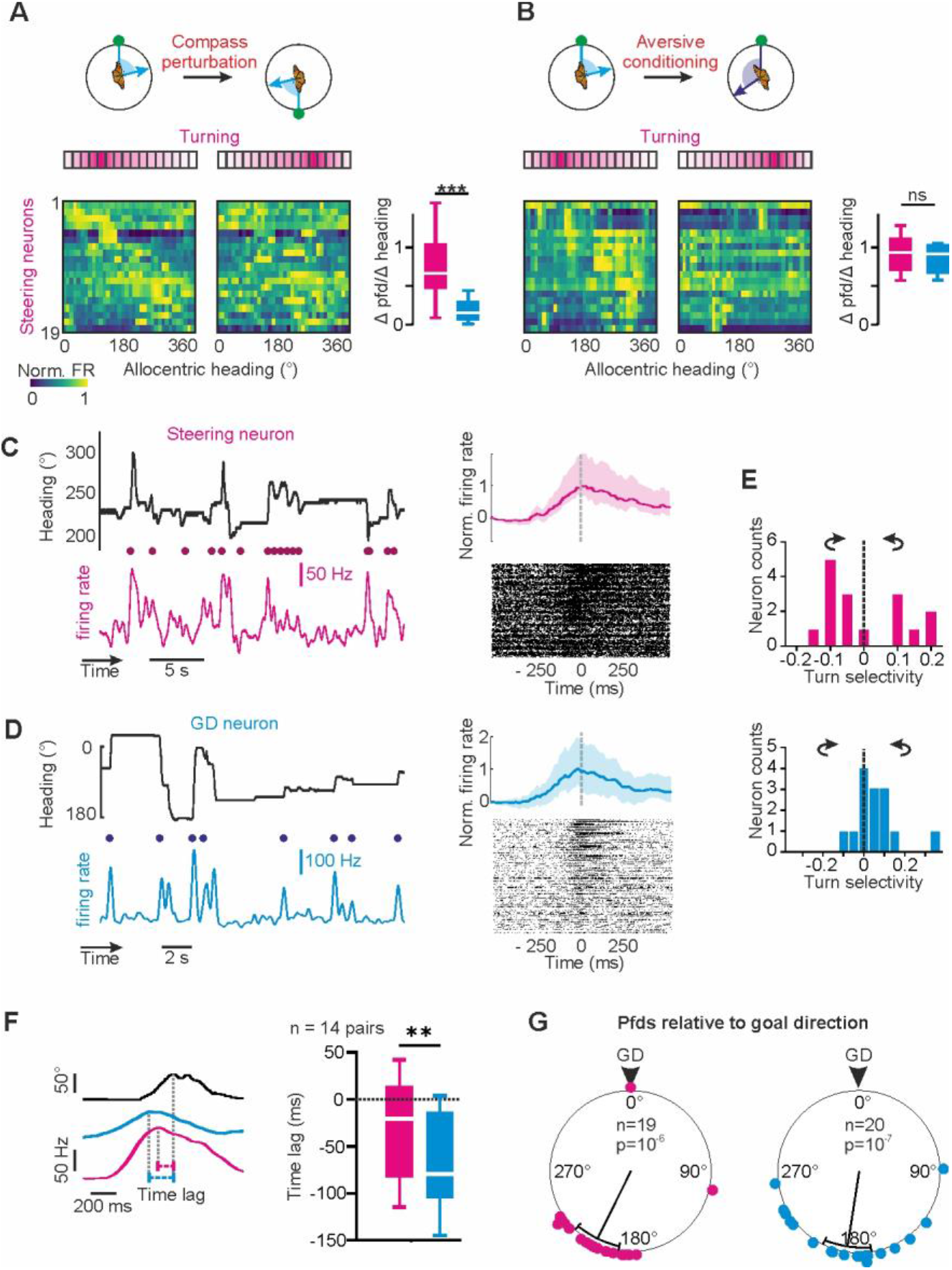
Turn coding in GD and steering neurons. **(A, B)** Change in angular tuning of steering neurons to compass perturbations (A) and aversive conditioning (B). Neurons were ordered according to their pfds before compass perturbations and conditioning. Right boxplots show the association between angular tuning and heading before and after compass perturbations (A) and conditioning (B) in steering and GD neurons. (**C, D**) Left: Example traces comparing heading (top) and neural firing rate (bottom) of a steering (C) and a GD neuron (D). Dots indicate time points of behavioral turns. Right: Sliding averages (top, shaded areas represent percentile) and raster plots (bottom) showing the firing rates of one steering (C) and one GD neuron (D) preceding turns (dashed line, time = 0). (**E**) Directional selectivity of steering (*top*, N = 16) and GD (*bottom*,N = 14) neurons. (**F**) Time lags, representing the duration of the neural activity preceding a turn, for pairs of simultaneously recorded GD (blue) and steering (magenta) neurons (dotted line indicates time point of behavioral turns). (**G**) Pfds of GD and steering neurons relative to the butterflies’ goal direction.

the suggested synaptic connection between GD and steering neurons (Matheson *et al*., 2022; Wystrach *et al*., 2020). In line with this proposition, pfds of GD (p < 0.001; V = 0.76; n = 20; V-test) and steering neurons (p < 0.001, V = 0.7, n = 19, V-test) were clustered in the direction opposite to the goal (Fig. 4G) which contrasts with the uniform distribution of pfds in HD neurons (p = 0.9; Z = 0.09; n = 13; Rayleigh test, fig. S13). Our results therefore suggest that the GD neurons closely interact with steering cells and activate them whenever the butterflies substantially deviate from their desired goal direction.

## DISCUSSION

We here discovered GD neurons in the insect central complex. The angular tuning of GD neurons changed when the butterfly’ s goal direction was reset (Fig. 2). More importantly the change was tightly associated with the change in goal direction (Fig. 3E). In contrast to this, compass perturbations did not affect the angular tuning in the very same neurons (Fig. 3). The tight association between neural tuning and goal-directed behavior *and* the robust selectivity for encoding the goal is compelling evidence that we have discovered the existence of GD neurons in invertebrates for the first time.

Our tetrode stainings (fig. S4) suggest that the insect GD neurons are localized in the fan-shaped body of the central complex, which is in line with recent hypotheses (Lu et al., 2022; Matheson *et al*.,2022; Stone *et al*., 2017; Wystrach *et al*., 2020). A network of GD neurons in monarch butterflies could represent the migratory southward direction as an activity bump across the 16 vertical columns of the fan-shaped body (Honkanen *et al*., 2019; Pisokas et al., 2022), similar to what has been demonstrated for the HD coding in the ellipsoid body (Hulse and Jayaraman, 2020; Seelig and Jayaraman, 2015). However, in contrast to the HD activity, the GD activity bump might not be yoked to the animal’s HD. Resetting the GD during aversive conditioning might have translocated the GD activity bump and hence the pfd of single GD neurons. We propose that a similar translocation of the GD activity bump might transform the butterfly’s southward migratory direction into a northward one (Guerra and Reppert, 2013).

Taken together, the navigation network of migratory butterflies consists of different neurons processing the current heading direction *and* goal direction, generating steering commands whenever the butterfly deviates from its course. In this study, we describe for the first time GD neurons in the insect brain and functionally discriminate them from HD and steering neurons. Despite being evolutionarily distant, our results show that the insect central complex houses similar GD neurons as the ones described in the mammalian brain, highlighting the computational power of the tiny insect brain in goal-directed navigation.

## Acknowledgments

We thank Marie Dacke, Lena van Giesen, Eric Warrant, Emily Baird, Kang Nian Yap, Alice Chou, and Stanley Heinze for their helpful comments on our manuscript. We thank Martin Strube-Bloss, Keram Pfeiffer, Wolfgang Rößler and Konrad Öchsner for their technical support. In addition, we thank Sergio Siles (butterflyfarm.co.cr) and Marie Gerlinde Blaese for providing us with monarch butterfly pupae.

## Funding

This work was funded by the Emmy Noether program of the German Research Foundation granted to BeJ (Grant number: EL784/1-1).

## Author contributions

Conceptualization (MJB, BeJ), Methodology and Formal Analysis

(MJB); Investigation (MJB, CK), Visualization (MJB, CK, BeJ), Funding acquisition (BeJ), Project administration and Supervision (BeJ), Writing - original draft (MJB, BeJ), Writing - review & editing (MJB, CK, BeJ).

## Competing interests

Authors declare that they have no competing interests.

## Data and materials availability

Matlab files with the calculated response parameters of the neurons together with the Matlab-scripts used for the analysis and Arduino scripts used for stimulus presentation are accessible from Datadryad: tba

## SUPPLEMENTS

## MATERIALS & METHODS

### Animals

Monarch butterflies (*Danaus plexippus*) were ordered as pupae from Costa Rica Entomological

Supply (butterflyfarm.co.cr) and kept in an incubator (HPP 110 and HPP 749, Memmert GmbH + Co. KG, Schwabach, Germany) at 25°C, 80% relative humidity and 12:12 light/dark-cycle conditions. After eclosion, the adult butterflies were transferred into another incubator (I-30VL, Percival Scientific, Perry, IA, USA) at 25°C and 12:12 light/dark condition. Adults had access to 15% sucrose solution *ad libitum*.

### Behavioral monitoring

A magnet (diameter = 3 mm; magnetic force = 4 N Supermagnete, Webcraft GmbH, Gottmadingen, Germany) was dorsally attached with dental wax (Article: 54895 Omnident, Rodgau Nieder-Roden, Germany) to the thorax of 32 butterflies. A second magnet at the end of a tungsten rod was used to connect the butterfly dorsally to an optical encoder (E4T miniature Optical Kit Encoder, US Digital, Vancouver, WA, USA) which measured the animal’s heading direction at a sampling rate of 100 Hz and at an angular resolution of 3°. Encoder signals were digitized (USB4 Encoder Data Acquisition USB Device, US Digital, Vancouver, WA, USA) and visualized in the US Digital software (USB1, USB4: US Digital, Vancouver, WA, USA). The optical encoder was vertically attached to a micro linear actuator (L12-R 50 mm 50:1 6 Volts, Actuonix Motion Devices, Saanichton, BC, Canada) that allowed us to control the butterfly’s suspension height using an Arduino MEGA 2560. The tethered butterfly could steer along any azimuth while being suspended at the center of a custom-built flight arena. The arena had an inner diameter of 32 cm and a height of 12 cm, and its upper inner circumference was equipped with 144 RGB-LEDs (Adafruit NeoPixel, Adafruit Industries, New York, New York, USA). The LED strip was mounted at an elevation of ~ 30° relative to the butterfly. One of these LEDs provided a single green light spot that served as a virtual sun stimulus (1.74 x 10^13^ photons/cm^2^/s and 1.2° angular extent at the butterfly’s eyes, as measured at the center of the arena). The angular position of the virtual sun was controlled by the Arduino MEGA 2560.

### Neural recordings

For neural recordings, one (N = 9) or three tetrodes (N = 23) were implanted in the butterfly central-complex. Each tetrode comprised a bundle of four 18 cm long and 12.5 μm thin copper wires (P155, Elektrisola, Reichshof-Eckenhagen, Germany) that were waxed tightly together. In experiments in which only one tetrode was implanted, the tetrode consisted of five copper wires (four recording and one differential wire). Tetrodes were carefully threaded through two Pebax^®^ tubes (each 2-4 cm in length; 0.026’ inner diameter; Zeus Inc, Orangeburg, SC, USA) that served as anchoring points to reversibly mount the tetrodes to a glass capillary. An additional copper wire served as grounding electrode and was immersed into the head capsule close to the butterfly’s neck. For aversive conditioning (N = 17), two stimulation copper wires (resistance ~10 MΩ) were waxed to the grounding electrode. All copper wires were soldered to gold pins and attached to an electrode interface board (EIB-18; Neuralynx Inc., Bozeman, MT, USA). In experiments in which three tetrodes were used, the tetrodes were fanned to maximally span 200-250 μm along the horizontal axis. Before each experiment, electrode resistances were measured with a nanoZ (Multi Channel Systems MCS GmbH, Reutlingen, Germany) and the electrode tips plated (Elektrolyt Gold solution, Conrad Electronic SE, Hirschau, Germany) to reduce the resistance of each electrode to ~0.1-1 MΩ. Tetrodes were reused for multiple experiments, after carefully trimming the tips and replating to the desired resistance.

Prior to obtaining neural signals of central-complex neurons, a monarch butterfly was horizontally restrained on a magnetic holder. To minimize movement artifacts during the recordings, the head was waxed to the thorax. The head capsule was opened dorsally and fat and trachea covering the brain surface were removed. To gain access to the central complex, the neural sheath on the dorsal brain surface was carefully removed using fine tweezers. The electrode bundle containing the grounding and the stimulation wires were inserted posteriorly in the head capsule, close to the butterfly’s neck. Tetrode tips were immersed in ALEXA 647 fluorophore coupled Hydrazide (A20502 diluted in 0.5 M KCl, Thermo Fisher Scientific GmbH, Dreieich, Germany) to quantify the tetrode position after each experiment. Recording tetrodes were then inserted into the brain, once per experiment. Tetrodes together with the glass capillary were attached to an electrode holder (M3301EH; WPI, Sarasota, FL, USA) and their positions controlled via a micromanipulator (Sensapex, Oulu, Finland). After adjusting the tetrode position along x- and y-axes, hemolymph fluid covering the brain was temporarily removed and the tetrodes were carefully moved along the z-axis to reach the central complex. While moving along the z-axis, band-pass filtered (600-6,000 Hz) neural signals were measured at a sampling frequency of 30 kHz. Neural signals were sent from the EIB-18 via an adapter board (ADPT-DUAL-HS-DRS; Neuralynx Inc., Bozeman, MT, USA) to a Neuralynx recording system (DL 4SX 32ch System, Neuralynx Inc., Bozeman, MT, USA). The neural activity was monitored using the software Cheetah (Neuralynx Inc., Bozeman, MT, USA). For setting a differential configuration, one electrode of the neighboring tetrode was set as a reference for the recording tetrode in the software. This means that the neural signals of each tetrode were referenced against the neural signal of an electrode of the neighboring tetrode. In cases in which only one tetrode was implanted, one of the five copper wires of the recording tetrode was set as a reference. To find visually sensitive neurons, the virtual sun was occasionally revolved clockwise and counterclockwise at an angular velocity of 60 deg/s around the insect’s head and the neural responses were visually quantified. After finding visually sensitive neurons at depths between 150-450 μm, the tetrode and the grounding wire were held in place by adding a two-component silicone elastomer (Kwik-Sil, WPI, Sarasota, FL, USA). After the Kwik-Sil hardened (~1 hour), the butterfly was carefully unrestrained and connected via the magnet to the end of the tungsten rod that was connected to the optical encoder. The tetrodes were carefully removed from the glass capillary and attached to a Pebax^®^ tube that was orthogonally oriented to the tungsten rod. To avoid wrapping the tetrode wires around the tungsten rod while the butterflies steered, the animals’ angular movements were restricted to 358°. To synchronize behavioral and neural recordings offline in Spike2 (version 9.0 Cambridge Electronic Devices, Cambridge, UK), a trigger signal was sent from the USB4 encoder via an ATLAS analog isolator (Neuralynx Inc., Bozeman, MT, USA) and the adapter board to the Neuralynx recording system at the onset of the behavioral recording. To temporally align stimulus presentations with the recorded neural activity, an analog output of the Arduino was sent via the ATLAS analog isolator to the Neuralynx recording system.

### Visualization of electrode tracks

After the neural recordings, the brain was dissected out of the head and fixated overnight in 4% formaldehyde at 4°C. The brain was then transferred into sodium-phosphate buffer and rinsed for 2 x 20 minutes in 0.1 M phosphate buffered saline (PBS) and 3 x 20 minutes in PBS with 0.3% Triton-X. The brain was dehydrated with an ascending ethanol series (30% - 100%, 15 minutes each) and immersed with a 1:1 ethanol-methyl-salicylate solution for 15 minutes, followed by a clearing step in methyl-salicylate for at least 1 hour. It was then embedded in Permount (Fisher Scientific GmbH, Schwerte, Germany) between two cover slips and scanned with a confocal microscope (Leica TCS SP2, Wetzlar, Germany) using a 20x water immersion objective (HC PL APO CS2 20x/0.75 IMM, Leica, Wetzlar, Germany). To visualize the tetrode position, we reconstructed the tetrode tracks in 3D using the software Amira 5.3.3 (ThermoFisher, Germany). To compare the tetrode positions from different experiments, we registered the tetrode position into the monarch butterfly standard central complex (Heinze *et al*., 2013). We used an affine (12-degrees of freedom), followed by an elastic registration to transfer the neuropils of the individual central complexes into the corresponding neuropils of the standard central complex. The registration and deformation parameters were then applied to the tetrode reconstruction to visualize the tetrodes in one frame of reference.

### Spike sorting and spike shape analysis

Neural recordings were spike sorted with the tetrode configuration implemented in Spike2 (version 9.00, Cambridge Electronic Devices, Cambridge, UK). We used four spike detection thresholds (two upper and two lower thresholds). The highest and lowest thresholds were set to avoid misclassifications of large voltage deflections occasionally arising from flight movements as spikes. The time window for template detection was set to 1.6 ms. After spike-sorting, a principal component analysis (PCA) was used to evaluate and to redefine spike clusters. Spike2 channels were exported as down-sampled Matlab files (3 kHz) and the remaining analysis was done with custom written scripts in MATLAB (Version R2021a, MathWorks, Natick, MA, USA). To analyze the spike shapes, the WaveMark channels containing the spike-waveforms were additionally exported as non-down-sampled Matlab files (30 kHz). For each neuron, spike-waveforms averaged from the first half of the experiment (compass perturbation) were correlated with the averaged spike-waveforms of the second half of the experiment (aversive conditioning) and statistically compared with the averaged spike-waveforms of the remaining neurons (Wilcoxon matched-pairs signed rank test: WSRT). This quantification allows us to statistically test whether neural recordings were stable throughout the experiment and assesses the quality of our spike-sorting analysis.

### Quantifying behavior and neural tuning

For behavioral analysis, we computed circular histograms by adding each data point of the optical encoder to the corresponding 10-degree heading bin. The animal’s preferred heading, represented by the mean vector, was computed with the CircStat toolbox for MATLAB. The flight directedness (*r*) was described with the mean vector strength which ranged between 0 (non-directed) to 1 (highly directed). Distributions of preferred headings of all animals were tested for uniformity with a Rayleigh test and visualized in Oriana (Version 4.01, Kovach Computing Services, Anglesey, Wales, UK).

Directional coding of neurons was quantified from circular plots. For each neuron and behavioral condition, i.e., sun position, pre-, post-conditioning, a circular plot was calculated that reflects the mean firing rate at different heading directions (10-degree bins). Circular statistics were then computed using the CircStat toolbox for MATLAB or in Oriana (Version 4.01, Kovach Computing Services, Anglesey, Wales, UK). First, angular sensitivity was determined by testing whether the mean firing rate deviated from a uniform distribution (Rayleigh test; significance level α = 0.05). If this was the case, we calculated the mean vector, or preferred firing direction (*pfd*), of a neuron.

### Dark experiments

To focus on neurons that showed an internal representation (GD neurons) or are tuned to idiothetic cues, i.e., in the absence of visual signals (HD neurons), we allowed the butterflies to orient on a Lab Jack prior to flight (Compact Lab Jack, Inc, Newton, New Jersey, USA). After the butterflies could steer in the presence of a virtual sun for a couple of minutes, we turned off the virtual sun and measured neural signals from the butterfly orienting in darkness. 113 out of 147 recorded neurons preserved their angular sensitivity when the butterflies were orienting in darkness and all subsequent neural analysis were based on these 113 neurons (Rayleigh test: significance level α = 0.05).

### Compass perturbation

To perturb the butterfly compass, we performed a similar experiment as the one performed in *Drosophila* (Green *et al*., 2019). However, instead of a vertical bar, we used the virtual sun as reference point of the insect compass. In the presence of the virtual sun, the butterfly flew for 9 min, and we changed the angular position of the sun every 90 s. In 15 experiments we changed the sun position in decreasing steps of 180°, 90°, 45°, 23°, and 15°. For the remaining 17 experiments, we exclusively changed the sun position in relatively large steps of 90° (3 times/experiment) or 180° (2 times/experiment). Preferred headings were measured every 90 s. Neural data were considered from three periods, in which the animals showed the highest flight directedness (*r*). Neurons were categorized regarding their changes in pfds in response to sun displacements. Hereby, we computed the circular variance of the heading offset (*CVH*) for each neuron. The heading offset represents the angular relation between pfd and behavioral heading directions. For heading-direction neurons, we suspect constant heading offsets throughout the experiment, i.e., the neurons’ pfds, should covary with the animal’s preferred heading. In addition to heading offset variance, we computed the circular variance of pfds (*CV*). This allowed us to measure the tuning stability. Both *CVH* and *CV* were weighted for each neuron by the following equation:

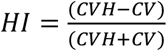

*HI* > 0 indicates that neural tuning can better be explained with a correlation to the animal’s heading (putative HD & steering neurons, n = 55), while *HI* < 0 indicate that neural tuning was unaffected by the animal’s heading and the sun’s position (putative GD neurons, n = 58). In addition, we correlated the binned neural response (10° bin size) measured prior to sun displacement with the one measured after displacement.

### Resetting the internal goal direction through aversive conditioning

To reset the butterfly’s internal goal direction without perturbing the compass system, we coupled the initial goal direction (± 90°) with electric shocks (*U* = 5 V; *I* = 0.5 μA). Prior to aversive conditioning (pre conditioning), the initial goal direction was visually determined by the experimenter while the butterfly oriented with respect to a static virtual sun. Depending on the butterfly’s flight directedness, this could take several minutes. To reset the goal direction by a significant amount, the butterfly received electric shocks whenever it flew in a sector containing the initial goal direction ± 90° (aversive conditioning). Electric shocks were controlled in the US Digital software (USB1, USB4: US Digital, Vancouver, WA, USA) that sent a signal from one of the USB4 output channels (USB4 Encoder Data Acquisition USB Device, US Digital, Vancouver, WA, USA) to the stimulus lines at the Neuralynx adapter board (ADPT-DUAL-HS-DRS; Neuralynx Inc., Bozeman, MT, USA). In parallel, the time course of stimulation was monitored by sending a digital signal from the USB4 to the Neuralynx system via the ATLAS analog isolator (Neuralynx Inc., Bozeman, MT, USA). Aversive conditioning took several minutes, depending on the butterfly’s performance. After aversive conditioning (post-conditioning), the butterfly was allowed to steer freely with respect to the virtual sun for several minutes. Note that the virtual sun’s azimuth was constant throughout the conditioning to avoid any compass perturbations. Heatmaps comparing the heading direction prior to and after conditioning were computed by normalizing the circular histograms containing the headings against the maximum bin. To roughly compare changes of preferred headings (behavior) and pfds (neurons) over time, we moved a sliding window in 10 s steps from the beginning of the aversive conditioning to the end of the experiment. To compute a preferred heading/pfd for each time window, it was necessary that the butterfly headed in each direction. Therefore, time window sizes were relatively large (mean/std: 262/78 s) and were set from the beginning of pre-conditioning to the time point of the first electric shock. Time courses of pfds were only used for the purposes of visualization. For quantitative analysis, we compared the angular tuning measured by circular plots between pre- and post-conditioning. Neurons that were categorized into putative HD/steering and GD neurons from the sun displacements were further categorized by calculating a GD index. The GD index is similarly calculated as the HD index, except that the difference in the goal offset was compared with the difference in pfd across the conditioning experiment. The goal offset describes the angular relation between the animal’s goal direction, i.e., preferred heading, and the neuron’s pfd. If this offset is similar after conditioning, then the neuron encodes the goal direction. In contrast to GD neurons, HD neurons should not change their angular tuning during conditioning and hence should have invariant pfds during conditioning. Based on the combination of HD and GD indices, we categorized four groups of neurons. 1) HD > 0; GD < 0; Neurons with constant heading offsets but varying goal offsets as one might predict for HD neurons (n = 13). 2) HD > 0; GD > 0; Neurons with constant heading and goal offsets receive compass and goal information as one might suspect from steering neurons (n = 19). 3) HD < 0; GD > 0; Neurons with varying heading offset but constant goal offset as one might suspect from GD neurons (n = 20). 4) HD < 0; GD < 0; Neurons with invariant pfds during sun displacement and conditioning and whose functions cannot be explicitly answered here (n = 13).

### Electric stimulation experiments in restrained butterflies

In control experiments aiming to test whether electric stimulation affects neural tuning, two stimulation copper wires (resistance: ~1 MΩ) were mounted on a single tetrode and inserted into the central complex of a restrained butterfly. The proximity of stimulation electrodes to the recording site allows one to undeniably test whether electric stimulation affects neural tuning in the central complex. For visual stimulation, the virtual sun was revolved clockwise and counterclockwise at an angular velocity of 60°/s around the butterfly. Electric stimulations were applied as pulses (1 ms) and repeated at 20 and 40 Hz with an electric current of 0.5-5 μA. Note that we even tested higher currents than the one used for aversive conditioning. Angular tuning, including pfds of 256 neurons were compared between pre and post stimulation (Wilcoxon matched-pairs signed rank test; WSRT).

### Testing for coding of turning behavior

To test for coding of flight turns, we determined the time points when the animal’s heading changed by more than 9°. We set 9° as turn threshold because the encoder’s angular resolution was 3° and deviations of ± 3° could represent variations in flight direction which may not represent substantial flight turns. In 1 s time windows, we examined the firing rate prior to (−500 ms) and after (+ 500 ms) the flight turns. Sliding averages of the neural activity were generated by applying a low-pass filter to the inter spike-intervals of the neurons. The neural activity in each time window was normalized to the firing rate 500 ms prior to the turn. Neural activity in time windows in which no flight turn occurred were considered as controls and statistically compared with the neural activity recorded during turns. Neurons were categorized as coding for flight turns if (i) modulations in the neural activity during flight turns were higher/lower than the modulations in neural activity during control (Wilcoxon p-test < 0.05) and (ii) if modulations in the neural activity during flight turns fitted a Gaussian distribution (> 0.7). Time lag between the peak firing rate and the maximum angular velocity (behavior) were computed by cross correlating the neural activity with the angular velocity. Negative time lags indicate that the neural activity changes prior to angular turns and vice versa. Neurons coding for flight turns were tested whether clockwise or counterclockwise turns elicited responses of different strengths by calculating a “turn selectivity”. Hereby, the peak firing rate in response to clockwise (*CW*) and counterclockwise (*CCW*) rotations were compared and weighted by the following formula:

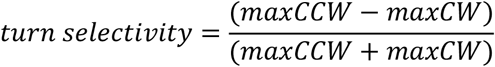

### Statistics

Circular statistics were performed in MATLAB and Oriana (Version 4.01, Kovach Computing Services, Anglesey, Wales, UK). All linear statistics were computed in GraphPad Prism 9 (GraphPad Software, San Diego, CA, USA). Sample sizes were not statistically pre-determined. Data distributions were tested for normality with a Shapiro-Wilk test. Normally distributed data were further analyzed with parametric statistical tests, while non-normally distributed data were tested with non-parametric tests. A Rayleigh test testing for uniformity of circular data was used to examine whether the flights were biased towards any direction. To statistically compare the angular tuning measured prior to and after compass perturbation across compass and putative GD neurons, we compared the correlation values obtained by correlating the angular tuning prior to sun displacement with the one measured after sun displacement with an unpaired t-test (Fig. 1I). Heading offsets and circular variances of pfds were statistically compared with a Mann-Whitney U test (MWU; Fig. 1J and fig. S6B). Variations in spike rate across compass and putative GD neurons were compared with a Mann-Whitney U test (Fig. S7). Changes in goal directions induced by aversive conditioning was statistically tested by comparing the distribution of GDs before conditioning (pre-conditioning) with the ones after conditioning (post conditioning) using a Mardia-Watson-Wheeler test (MWW) (fig. S8B). Flight directedness prior to and after conditioning was compared with a paired t-test (fig. S8C). To compare the tuning stability prior to compass perturbation and aversive conditioning with the one measured after compass perturbation and aversive conditioning, we statistically compared the correlation values obtained by comparing the angular tunings with an ordinary one-way ANOVA across different neuron types, i.e., HD, GD, and steering neurons (fig S12B). Note when comparing between two neuron types, we used a Mann-Whitney U test (Fig. 3D). A Mann-Whitney U test was used to statistically compare the changes in pfds induced by aversive conditioning in GD neurons (Fig. 3C) and when comparing pfd changes induced by compass perturbation and aversive conditioning between GD and steering neurons (Fig. 4A and 4B). Time lags of turn coding were statistically compared across steering and GD neurons with a Mann-Whitney U test (Fig. 4F). Hereby, only pairs (n = 14 pairs) of simultaneously recorded steering and GD neurons were considered because a comparison of time lags across different experiments were unprecise due to the relatively low sampling rate of the optical encoder. The consistency of goal offsets for putative GD, and HD neurons across the conditioning was statistically compared with Mann-Whitney U test (Fig. 4E). Later, the putative GD neurons were divided into GD and steering neurons and their goal offset stability across conditioning compared with the one of HD neurons (Kruskal-Wallis test; One-Way ANOVA; Fig. S12C). With a Rayleigh test, we examined whether pfds of HD neurons were uniformly distributed (fig. S13) and a V-test (expected 180°) allowed us to demonstrate that pfds of GD and steering neurons were clustered at 180° (Fig. 4G).

Statistical tests were always two-sided. Data collection and analysis were not conducted blind to the conditions of the experiments. For neural recordings, stimulus presentation was pseudorandomized. We excluded 34 of the 147 recorded neurons, because of the lack of angular tuning when the butterflies oriented in darkness on a platform prior to flight (Rayleigh test: p > 0.05; see also fig. S9).

### Data and codes

Matlab files with the calculated response parameters of the neurons together with the Matlab-scripts used for the analysis and Arduino scripts used for stimulus presentation are accessible from Datadryad: https://tba

---

**Figure S1.**
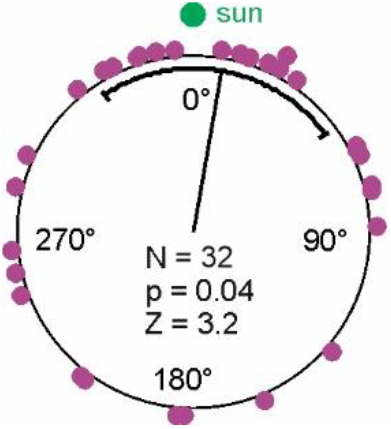
Distribution of preferred headings relative to the virtual sun. Circularplot visualizing preferred headings of 32 butterflies, sun positioned at 0°. Statistics from a Rayleigh test, testing against a uniform distribution is depicted in the center of the circularplot.

**Figure S2.**
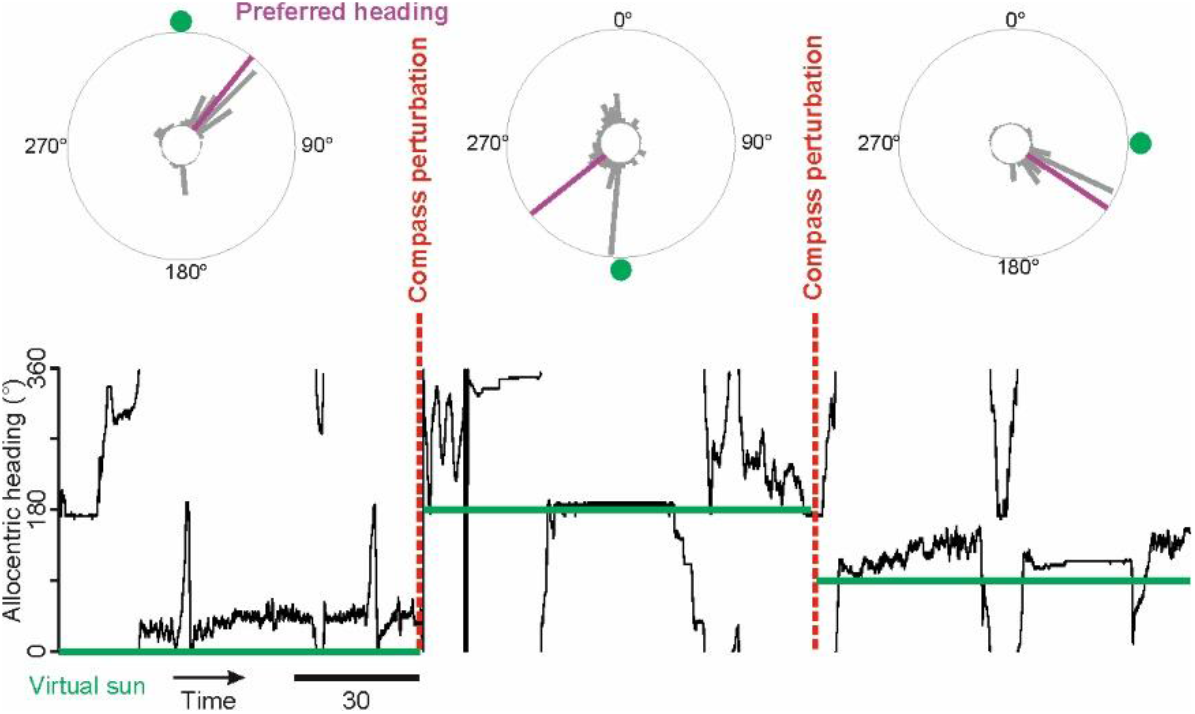
Behavioral performance of a butterfly whose compass polarity was changed by displacing the virtual sun. Butterfly’s heading (*black*) as a function of time. The angular position of the virtual sun is depicted in *green*. Every 90 seconds the virtual sun was displaced (compass perturbation). Circular histograms demonstrate the butterfly’s heading at different virtual sun positions. Note that the butterfly changed its preferred heading to set a consistent goal heading relative to the virtual sun.

**Figure S3.**
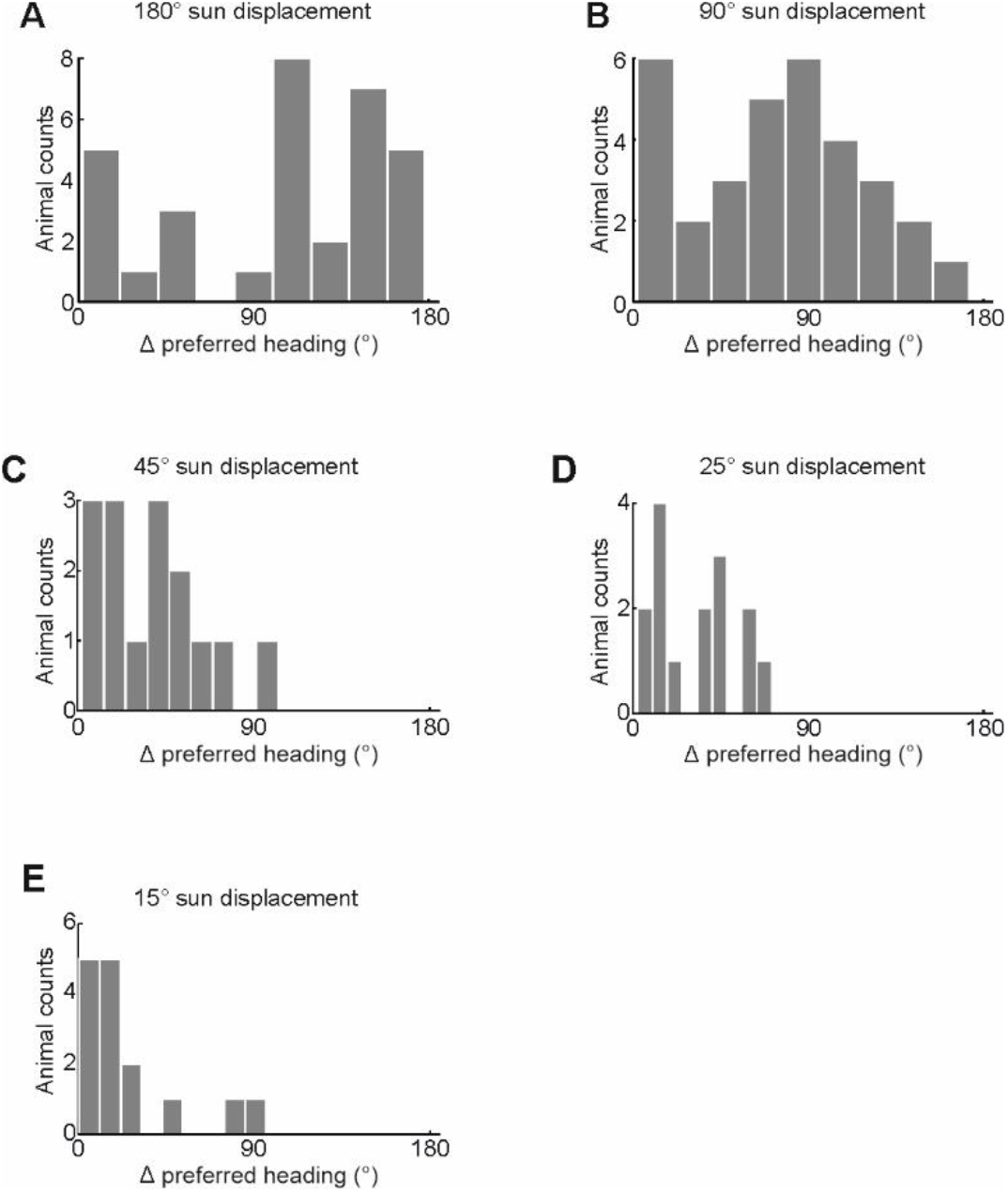
Summary of heading changes induced by displacing the virtual sun at different angular positions. Histograms showing the butterflies’ change of heading after 180° (**A**), 90° (**B**),45° (**C**), 25° (**D**), and 15° (**E**) sun displacements.

**Figure S4.**
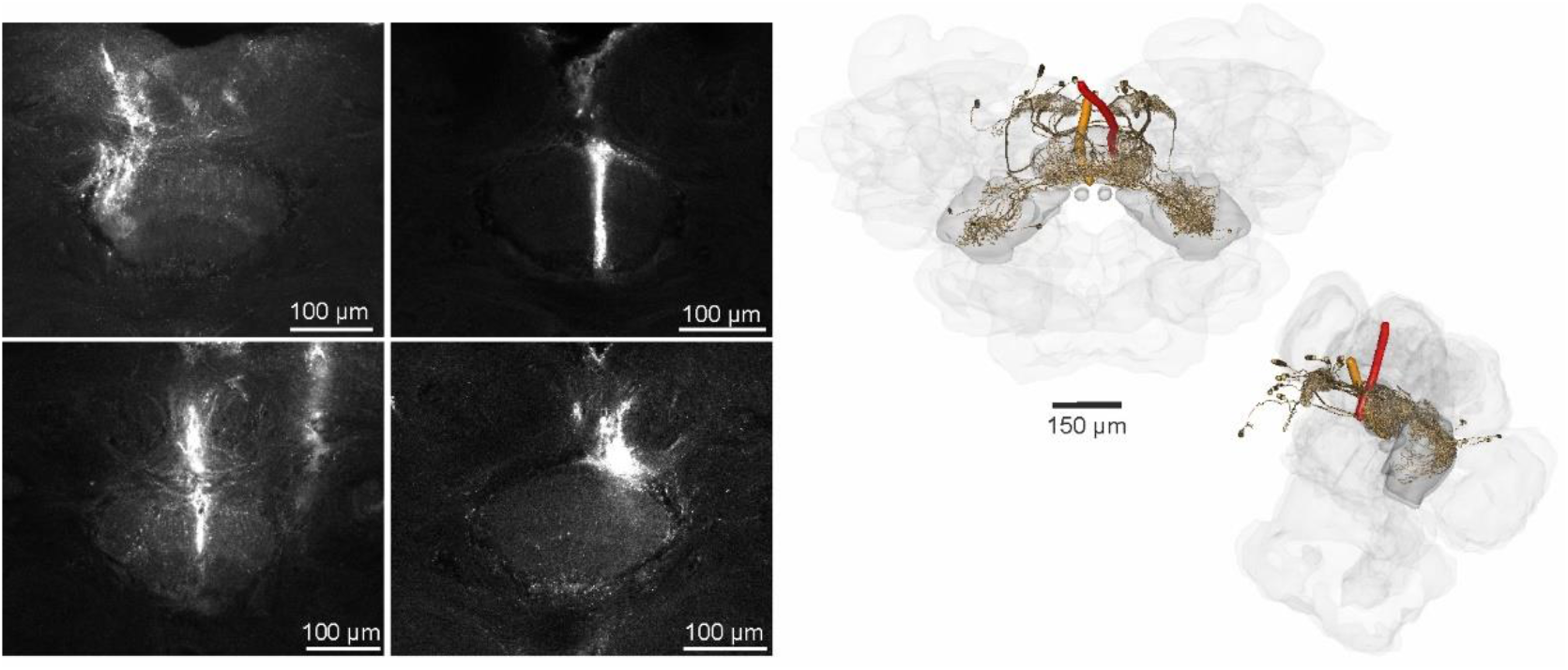
Visualiztion of tetrode positions in the butterfly brain. *Left:* Z-stacks from the fan-shaped body of the central complex showing stainings from four example tetrode tracks. *Right:* Anterodorsal (*left*) and lateral (*right*) view of 3D reconstructed tetrode tracks that correspond to the example neurons presented in Fig.1G and 2F. Prominent central-complex neurons (*brown*) are visualized.

**Figure S5.**
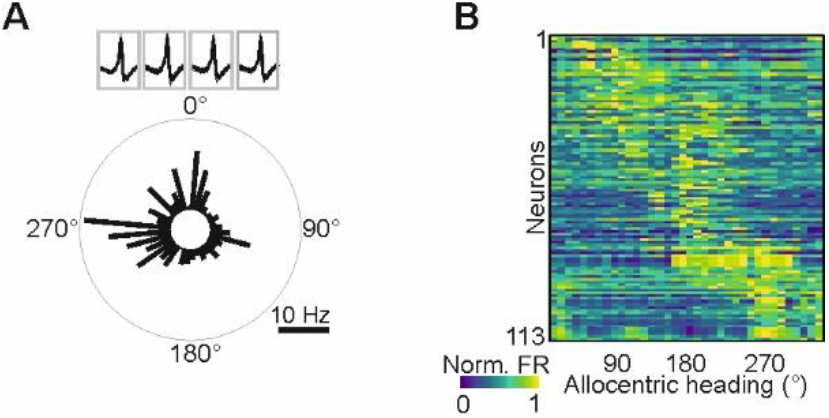
Central-complex neurons with an internal representation of directions. Angular tuning of an example (**A**) and 113 (**B**) neurons measured when the butterfly actively rotated on a platform in darkness. All neurons were spatially tuned according to a Rayleigh test (p < 0.05). Neurons are ordered according to their preferred firing directions.

**Figure S6.**
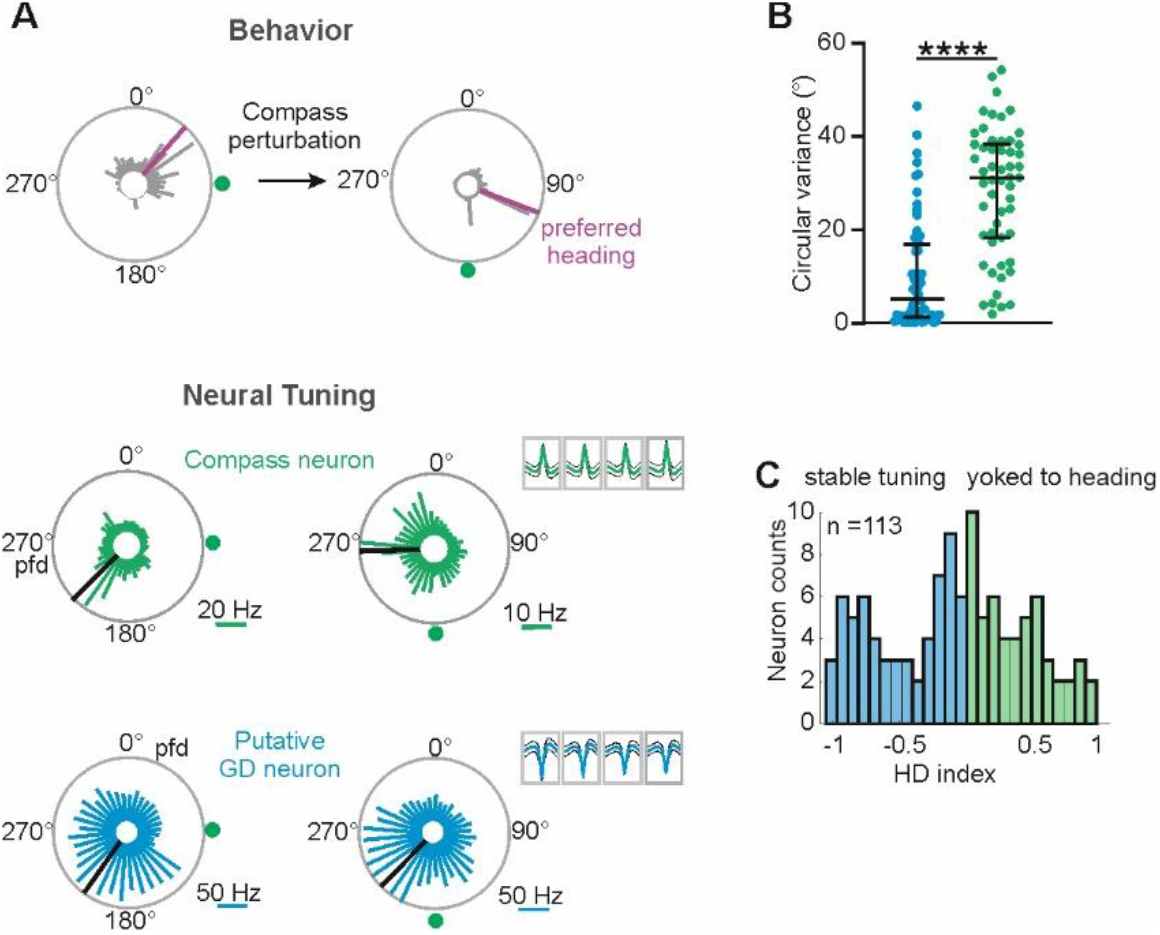
Discrimination of compass and putative GD neurons after sun displacements. (**A**) Behavioral response (upper circular plots) and neural tuning of two example neurons (green and blue) in response to a 90° sun displacement. Respectively, the preferred heading and the preferred firing direction (pfd) is indicated by purple and black bars. The mean and percentile of the spike wave forms from each tetrode electrode is depicted in the right upper corner of the circular plots. Note that the angular tuning of the green neuron follows the behavioral response and the pfd changes by about 90° while the angular tuning of the blue neuron is invariant. (**B**) Comparison of circular variances of pfds in response to compass perturbations for putative GD (*blue*, n = 58) and compass (*green*, n = 55) neurons. (**C**) Histogram of measured HD indices. Indices indicate that the angular tuning is stable (negative) or associated with the butterfly’s preferred heading (positive).

**Figure S7.**
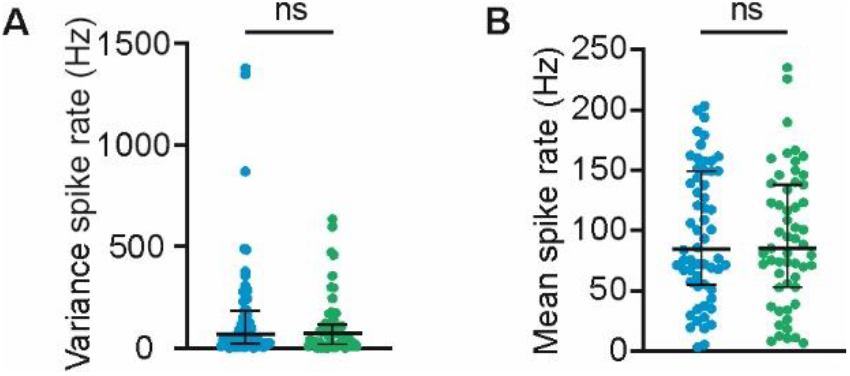
Overview of spike rate parameters for GD (*blue*) and compass neurons (*green*) during compass perturbations. (**A**) Spike rate variance for GD and compass neurons (Mann Whitney U test, variance: p = 0.59, U = 1501, n = 55 compass neurons; n = 58 GD neurons). (**B**) Mean spike rate for GD and compass neurons (Mann Whitney U test, variance: p = 0.75, U = 1540, n = 55 compass neurons; n = 58 GD neurons).

**Figure S8.**
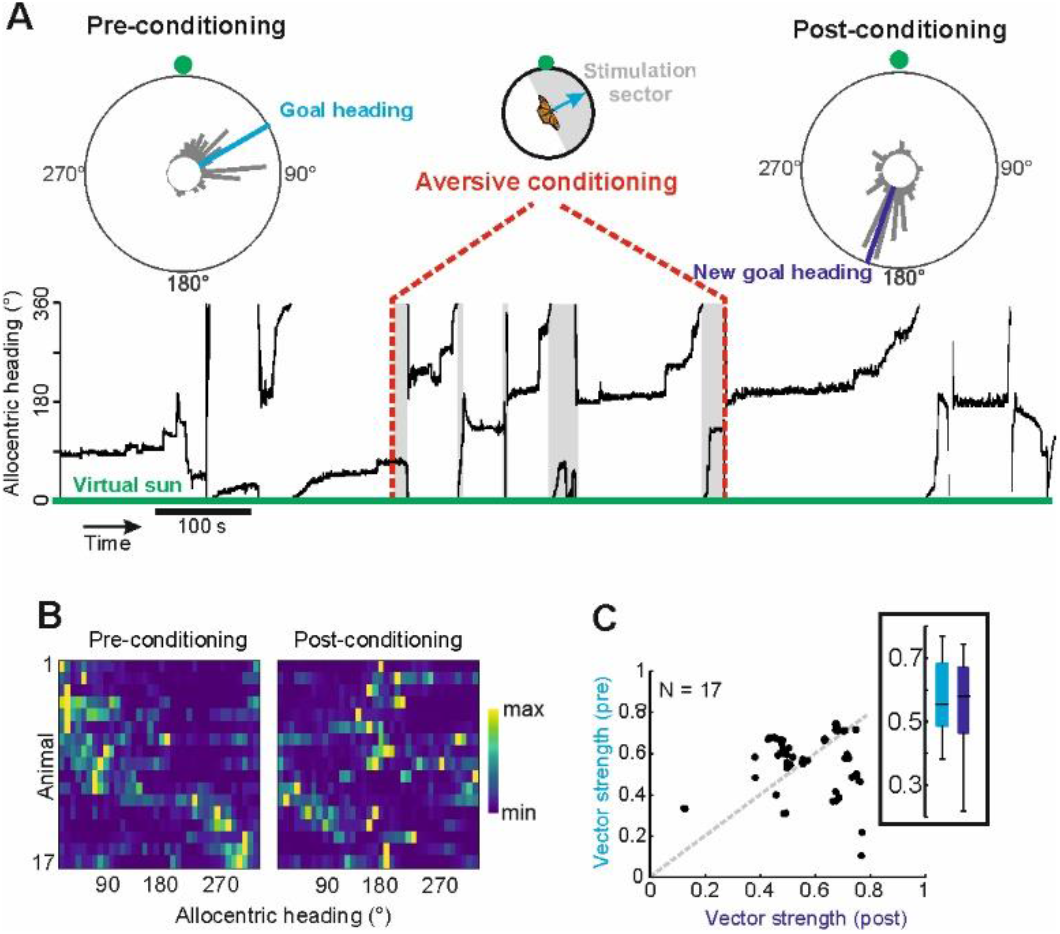
Behavioral performance of butterflies in response to aversive conditioning. (**A**) Heading plotted as a function of time. Gray boxes highlight periods of electric shocks. The virtual sun was held in place at 0°. Circular plots summarize the heading before and after conditioning. (**B**) Distribution of normalized heading of 17 butterflies before (*left heatmap*) and after (*right heatmap*) conditioning. Butterflies were ordered according to their initial GD at pre-conditioning. (**C**) Flight directedness, represented by the vector strength, was unaffected by conditioning.

**Figure S9.**
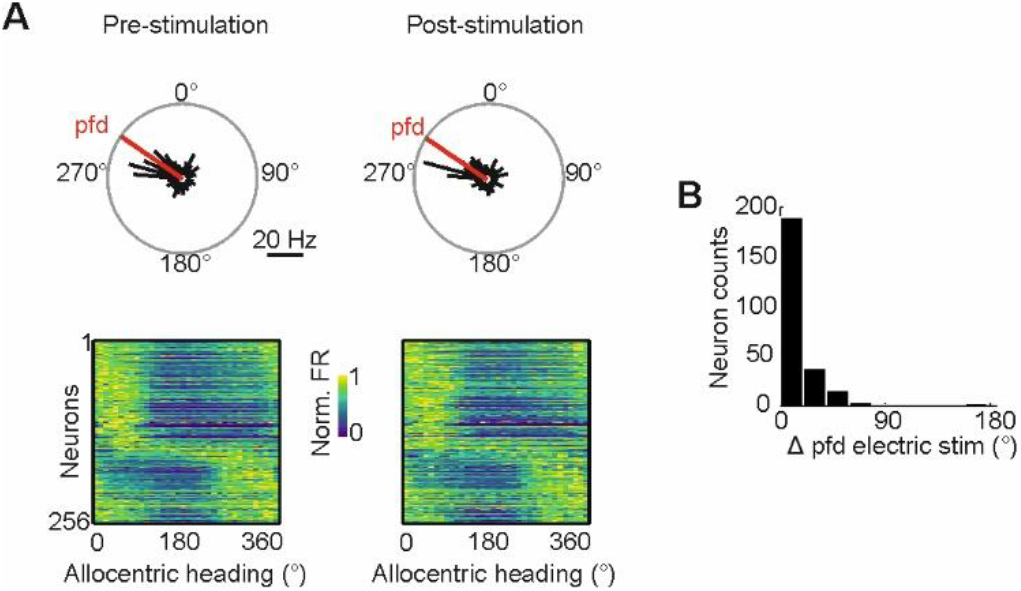
Electric stimulation in the central complex does not affect neural tuning. (**A**) *Upper:* Angular tuning of a central-complex neuron measured with the virtual sun revolving around a restrained butterfly before (Pre-stimulation) and after (Post-stimulation) electric stimulation. Red line represents the preferred firing direction (pfd). *Lower*: Angular tuning of 256 central-complex neurons measured before and after electric stimulation. Neurons are ordered according to their pfd. (**B**) Electric stimulation in the central complex does not change the neurons’ pfds (Wilcoxon matched-pairs signed rank test, p = 0.63, W = 1136, n = 256).

**Figure S10.**
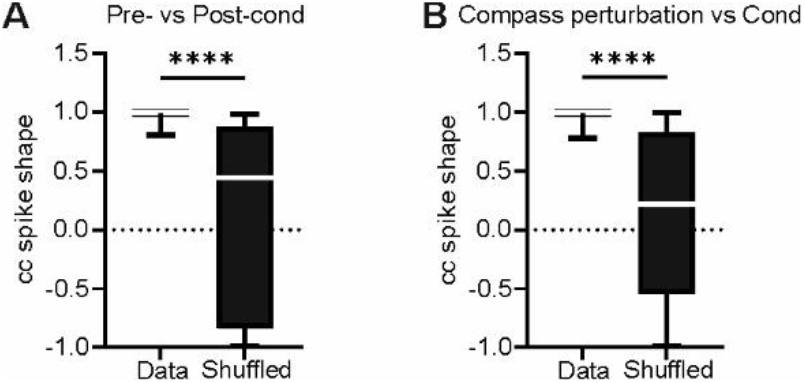
Spike shape stability of neurons recorded during the experiment. Correlation values of spike shapes measured before and after conditioning (**A**) and during compass perturbation and conditioning (**B**) were compared with correlation of spike shapes shuffled across randomly selected neurons [Wilcoxon matched-pairs signed rank test, p < 10^-5^, W = −3395 (A), p < 10^-5^, n = 82; W = −10276, n = 144 (B)].

**Fig. S11.**
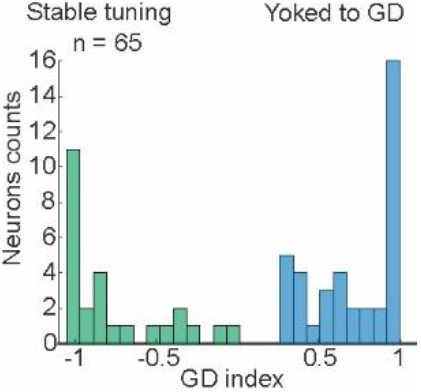
Distribution of goal direction indices from 65 neurons comparing stable tuning against tuning yoked to the goal direction. Histogram of measured goal direction indices (GD indices). Indices indicate that the angular tuning is stable (negative) or yoked to the butterfly’s goal direction (positive).

**Figure S12.**
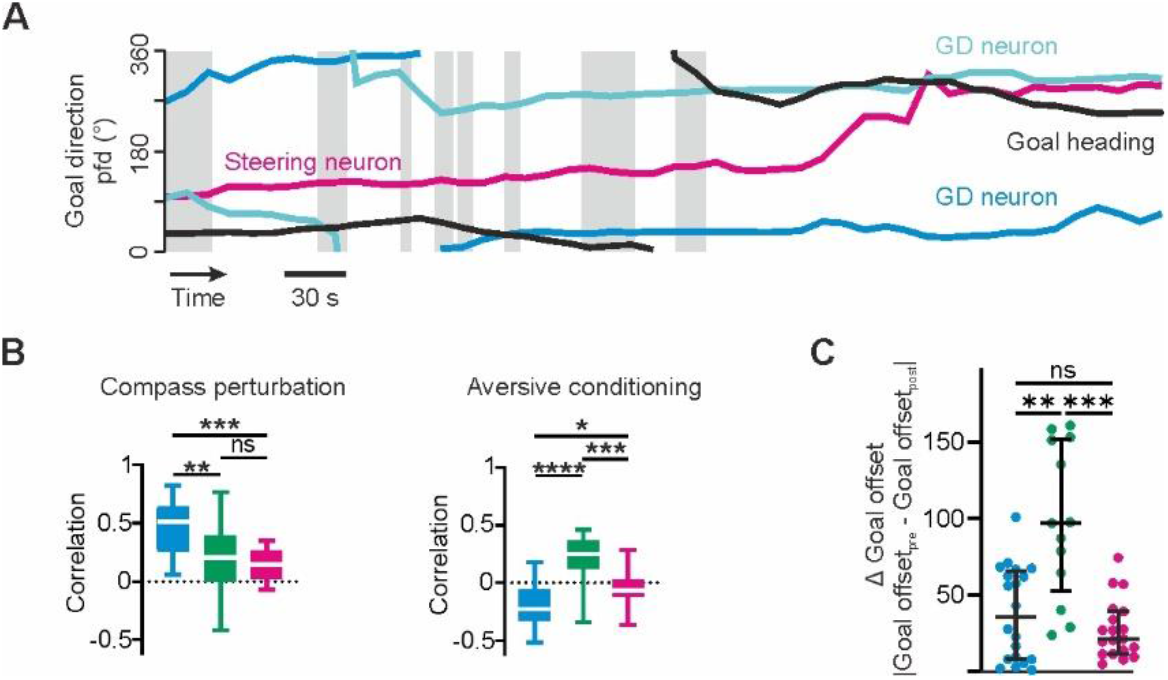
Angular tuning changes of GD, HD, and steering neurons in response to aversive conditioning. (**A**) Goal heading (black line) and preferred firing directions (pfds; colored lines) of example neurons (magenta: steering neuron, blue: GD neurons) plotted as a function of time (Time = 0 start of conditioning). Gray boxes highlight periods of electric stimulation. (**B**) Correlation of angular tuning before and after compass perturbations (*left*) or conditioning (*right*). (**C**) Differences of goal offsets prior to and after conditioning. The lower the goal offset differences the more was the pfd yoked to the goal direction. Note that angular tuning of both GD and steering neurons was dependent on the butterfly’s goal direction while the angular tuning of HD neurons was independent from the goal direction.

**Figure S13.**
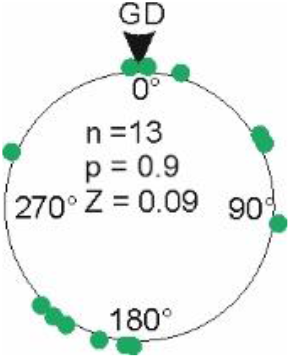
Distribution of pfds of HD neurons relative to the butterflies’ GDs.

## Notes

### Competing Interest Statement

The authors have declared no competing interest.

## REFERENCES

Beetz, M.J., Kraus, C., Franzke, M., Dreyer, D., Strube-Bloss, M.F., Rössler, W., Warrant, E.J., Merlin, C., and el Jundi, B. (2022). Flight-induced compass representation in the monarch butterfly heading network. Curr Biol 32, 338–+. 10.1016/j.cub.2021.11.009.

Ben-Yishay, E., Krivoruchko, K., Ron, S., Ulanovsky, N., Derdikman, D., and Gutfreund, Y. (2021). Directional tuning in the hippocampal formation of birds. Curr Biol. 10.1016/j.cub.2021.04.029.

Dacke, M., and el Jundi, B. (2018). The dung beetle compass. Curr Biol 28, R993–R997. 10.1016/j.cub.2018.04.052.

Geva-Sagiv, M., Las, L., Yovel, Y., and Ulanovsky, N. (2015). Spatial cognition in bats and rats: from sensory acquisition to multiscale maps and navigation. Nat Rev Neurosci 16, 94–108. 10.1038/nrn3888.

Green, J., Vijayan, V., Mussells Pires, P., Adachi, A., and Maimon, G. (2019). A neural heading estimate is compared with an internal goal to guide oriented navigation. Nat Neurosci 22, 1460–1468. 10.1038/s41593-019-0444-x.

Guerra, P.A., and Reppert, S.M. (2013). Coldness Triggers Northward Flight in Remigrant Monarch Butterflies. Current Biology 23, 419–423. 10.1016/j.cub.2013.01.052.

Heinze, S., Florman, J., Asokaraj, S., El Jundi, B., and Reppert, S.M. (2013). Anatomical basis of sun compass navigation II: the neuronal composition of the central complex of the monarch butterfly. J Comp Neurol 521, 267–298. 10.1002/cne.23214.

Heinze, S., and Reppert, S.M. (2011). Sun Compass Integration of Skylight Cues in Migratory Monarch Butterflies. Neuron 69, 345–358. 10.1016/j.neuron.2010.12.025.

Honkanen, A., Adden, A., da Silva Freitas, J., and Heinze, S. (2019). The insect central complex and the neural basis of navigational strategies. J Exp Biol 222. 10.1242/jeb.188854.

Hulse, B.K., and Jayaraman, V. (2020). Mechanisms Underlying the Neural Computation of Head Direction. Annu Rev Neurosci 43, 31–54. 10.1146/annurev-neuro-072116-031516.

Lu, J., Behbahani, A.H., Hamburg, L., Westeinde, E.A., Dawson, P.M., Lyu, C., Maimon, G., Dickinson, M.H., Druckmann, S., and Wilson, R.I. (2022). Transforming representations of movement from body-to world-centric space. Nature 601, 98–104. 10.1038/s41586-021-04191-x.

Martin, J.P., Guo, P.Y., Mu, L.Y., Harley, C.M., and Ritzmann, R.E. (2015). Central-Complex Control of Movement in the Freely Walking Cockroach. Curr Biol 25, 2795–2803. 10.1016/j.cub.2015.09.044.

Matheson, A.M.M., Lanz, A.J., Medina, A.M., Licata, A.M., Currier, T.A., Syed, M.H., and Nagel, K.I. (2022). A neural circuit for wind-guided olfactory navigation. Nat Commun 13, 4613. 10.1038/s41467-022-32247-7.

Mouritsen, H., and Frost, B.J. (2002). Virtual migration in tethered flying monarch butterflies reveals their orientation mechanisms. Proc Natl Acad Sci U S A 99, 10162–10166. 10.1073/pnas.152137299.

Nyberg, N., Duvelle, E., Barry, C., and Spiers, H.J. (2022). Spatial goal coding in the hippocampal formation. Neuron 110, 394–422. 10.1016/j.neuron.2021.12.012.

Petrucco, L., Lavian, H., Wu, Y.K., Svara, F., Štih, V., and Portugues, R. (2022). Neural dynamics and architecture of the heading direction circuit in a vertebrate brain. bioRxiv, 2022.2004.2027.489672. 10.1101/2022.04.27.489672.

Pisokas, I., Rossler, W., Webb, B., Zeil, J., and Narendra, A. (2022). Anesthesia disrupts distance, but not direction, of path integration memory. Current Biology 32, 445–+. 10.1016/j.cub.2021.11.039.

Sarel, A., Finkelstein, A., Las, L., and Ulanovsky, N. (2017). Vectorial representation of spatial goals in the hippocampus of bats. Science 355, 176–180. 10.1126/science.aak9589.

Seelig, J.D., and Jayaraman, V. (2015). Neural dynamics for landmark orientation and angular path integration. Nature 521, 186–+. 10.1038/nature14446.

Stone, T., Webb, B., Adden, A., Ben Weddig, N., Honkanen, A., Templin, R., Wcislo, W., Scimeca, L., Warrant, E., and Heinze, S. (2017). An Anatomically Constrained Model for Path Integration in the Bee Brain. Curr Biol 27, 3069–+. 10.1016/j.cub.2017.08.052.

Takahashi, S., Hombe, T., Matsumoto, S., Ide, K., and Yoda, K. (2022). Head direction cells in a migratory bird prefer north. Sci Adv 8, eabl6848. 10.1126/sciadv.abl6848.

Taube, J.S., Muller, R.U., and Ranck, J.B., Jr. (1990). Head-direction cells recorded from the postsubiculum in freely moving rats. I. Description and quantitative analysis. J Neurosci 10, 420–435.

Varga, A.G., and Ritzmann, R.E. (2016). Cellular Basis of Head Direction and Contextual Cues in the Insect Brain. Curr Biol 26, 1816–1828. 10.1016/j.cub.2016.05.037.

Vinepinsky, E., Cohen, L., Perchik, S., Ben-Shahar, O., Donchin, O., and Segev, R. (2020).

Representation of edges, head direction, and swimming kinematics in the brain of freely-navigating fish. Sci Rep 10, 14762. 10.1038/s41598-020-71217-1.

Wan, G., Hayden, A.N., Iiams, S.E., and Merlin, C. (2021). Cryptochrome 1 mediates light-dependent inclination magnetosensing in monarch butterflies. Nat Commun 12, 771. 10.1038/s41467-021-21002-z.

Wystrach, A., Le Moёl, F., Clement, L., and Schwarz, S. (2020). A lateralised design for the interaction of visual memories and heading representations in navigating ants. bioRxiv, 2020.2008.2013.249193. 10.1101/2020.08.13.249193.

